# Decoding the Plasma Proteomic Landscape of Clear Cell Renal Cell Carcinoma Reveals Diagnostic and Prognostic Liquid Biopsy Biomarkers

**DOI:** 10.64898/2026.08.03.742031

**Authors:** Harini Lakshminarayanan, Dorothea Rutishauser, Peter Schraml, Daniel Eberli, Hella Bolck, Holger Moch

## Abstract

Clear cell renal cell carcinoma (ccRCC) remains the most lethal urological malignancy, with high metastatic rates, both at initial diagnosis and during disease progression, contributing to poor survival outcomes. Current diagnostic and prognostic approaches rely primarily on histopathology, limiting early detection of localized disease and relevant intervention for metastatic patients. Here, we performed the most extensive to-date mass spectrometry-based discovery profiling of longitudinal plasma samples collected across multiple clinical follow-up points spanning up to five years post-diagnosis., to characterize the circulating plasma proteome and identify biomarkers for localized and metastatic disease. Network analysis identified protein modules enriched in pathways involved in matrix remodeling and metabolic deregulation, perpetuating the ccRCC phenotype.

A five-protein signature, comprising PRL, THBS1, ANGPT1, IGFBP1, and SRGN, demonstrated high diagnostic performance for localized ccRCC. Notably, PRL appeared as a promising stand-alone biomarker (AUC = 0.812), with independent validation confirming its utility as a diagnostic biomarker. Importantly, a six-protein signature (AMBP, C1S, C2, IGFBP3, RASGRP2, TFRC) stringently distinguished metastatic from high-grade non-metastatic ccRCC cases. Further validation of these signatures could inform clinical decision-making, enabling early detection of metastasis and minimal residual disease and real-time longitudinal monitoring for ccRCC patients.

**Statement of Significance:** This study presents the most comprehensive longitudinal plasma proteomic dataset for ccRCC to date, defining robust circulating protein biomarker signatures for both localized and metastatic disease, and establishing a proteomic landscape for minimally invasive, real-time monitoring and improved clinical management of ccRCC patients.

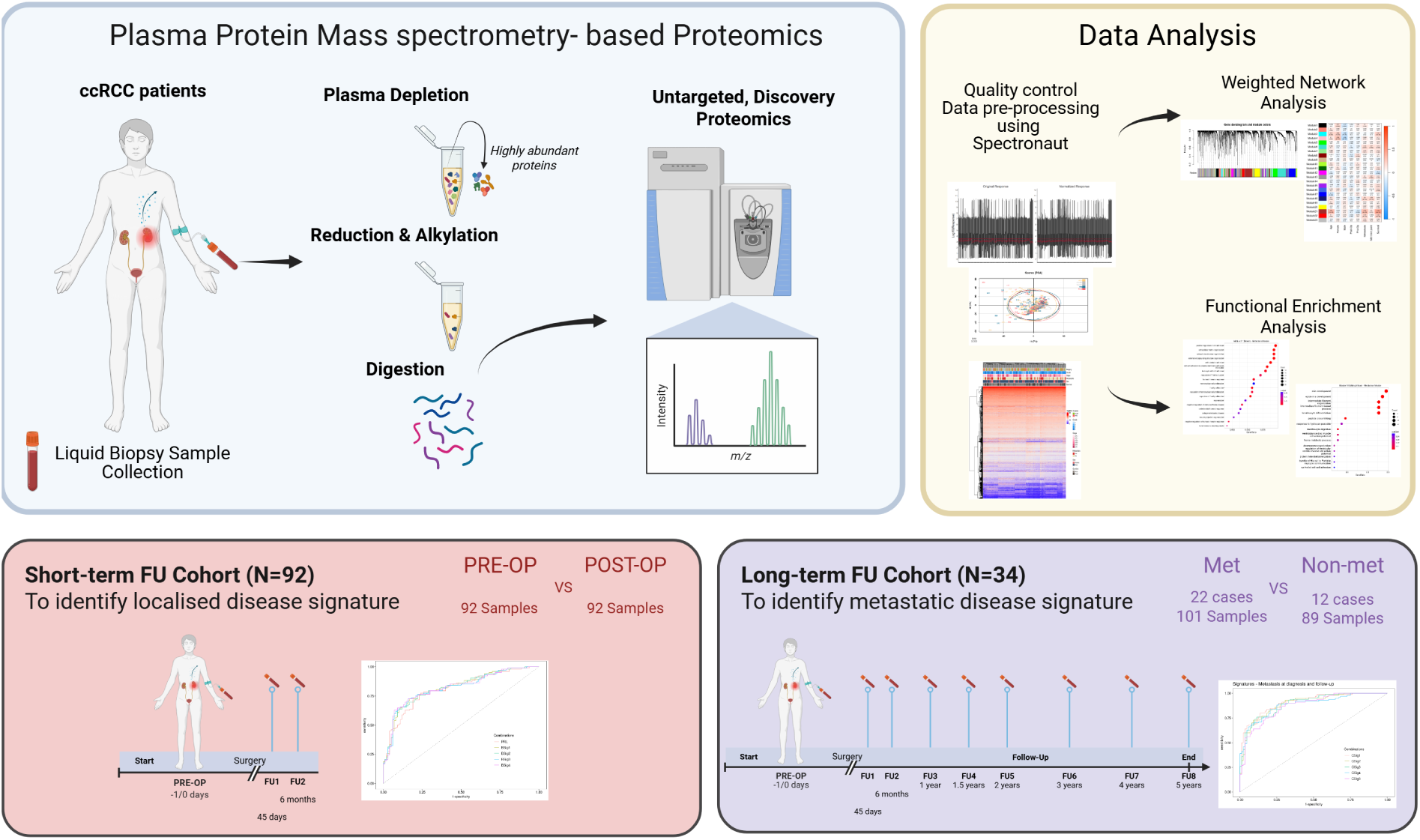

## Introduction

Clear cell renal cell carcinoma (ccRCC) is the most common renal cancer subtype, accounting for 80% of RCC in adults [1, 2]. Notably, around 30% of ccRCC patients present with metastasis at time of initial diagnosis (synchronous metastasis), and another 30% develop metastatic disease (late metastasis) as the cancer progresses [3]. This broad variation in stages of detection presents the challenge of a wide range of clinical outcomes. Survival rates vary from 6 months to 5 years, where mRCC patients face abysmal prognosis with overall survival of 10% as compared to a higher survival rate of 85% for localized disease. These differing clinical outcomes are further influenced by several distinct disrupted molecular mechanisms and high inter-tumor heterogeneity in ccRCC, stemming from chromosomal complexities and clonal evolution [4, 5]. *VHL* inactivation due to the landmark loss of the short arm of chromosome 3p plays a central role in the etiology of ccRCC. This results in strikingly disrupted hypoxic signaling and aberrant cellular metabolism following the activation of VHL-HIF signaling axis, driving disease progression [6–8].

Although the tumor molecular landscape is well studied, the translation of biomarkers into clinical practice has been limited. Current clinical guidelines for diagnosis require the histopathological characterization of tumor tissue sections, with sufficient categorization of the tumor grade, stage and differentiation features. This is further used for patient risk stratification for prognosis [2]. The International Metastatic Renal Cell Carcinoma Database Consortium’s (IMDC) score is the prognostic model used in clinics, taking into account six factors, namely Karnofsky performance status (KPS) (categorizing the patient’s functional impairment), time from diagnosis to first-line targeted therapy, hemoglobin concentration, neutrophil count, platelet count, and serum calcium concentration. While this model has been validated [9, 10], it has limited utility in the treatment landscape lacking ccRCC-specific molecular information. Simultaneous to the currently varied survival outcomes, ccRCC is also increasingly incidentally detected during unrelated abdominal imaging, due to it’s asymptomatic initial progression [1]. Therefore, reliable molecular biomarkers for diagnosis and prognosis are an urgent clinical need, with longitudinal monitoring of patients imperative to identify risk of disease dissemination and metastasis, detect minimal residual disease, and support informed clinical decision-making.

Minimally invasive technologies like liquid biopsy present the opportunity to monitor patients, across time, by analyzing small volumes of their biological fluids including blood. Investigating the plasma circulome of cancer patients, such as circulating nucleic acids, extracellular vesicles, and proteins, could allow the early detection of dissemination and metastasis [11]. Among these biomolecules, proteins directly affect physiology and pathology. Plasma proteins are likely to reveal organ- and tumor-specific information in circulation [12] and can capture molecular changes associated with both primary tumors and metastatic sites, providing a window to disease dynamics [13]. Previously, single protein candidates have been selected and tested in RCC patient blood, based on known RCC-specific molecular alterations. Markers implicated in the VHL-HIF signaling axis, such as KIM1 [14], CAIX [15], HIG2 [16], and IMP3 [17] have previously been tested using immunoassays, in either patient plasma or urine samples. However, these markers were not studied in a longitudinal setting and have not since demonstrated clinical utility for diagnosis or prognostication, emphasizing the need for integrating multiple biomarkers into a panel, to ensure high sensitivity and specificity. Targeted affinity-based techniques using oncology-related protein panels have noted potential proteins as capable of distinguishing disease from healthy control [18], however such studies are limited by the choice of the target panel, preventing large discovery studies. In the meantime, mass spectrometry-based proteomics studies in ccRCC have also been conducted, but these were still limited, either by small cohorts without follow-up samples or the challenges arising from the large dynamic range of proteins in human plasma [19–21]. Plasma protein biomarkers are indeed technically challenging to identify due to the complexity of blood as a disease medium. Blood biomarker discovery is marred by 12 highly abundant proteins contributing to 95% of the total plasma proteome [22] and several strategies, including depletion, are being defined to access the potential biomarker-rich low-abundance region in plasma [23]. The successful integration of these depletion techniques with high-sensitivity, high-coverage quantification through parallel mass spectrometry [24] provides a valuable opportunity to characterize ccRCC plasma and identify protein signatures for localized and metastatic disease.

In our study, we apply mass spectrometry-based proteomic profiling to an exceptionally large, rigorously designed ccRCC cohort, incorporating longitudinal plasma samples collected across several years. This unique dataset enables a comprehensive characterization of the circulating proteomic landscape supporting the clinical progression of ccRCC. Using this data, we delineate the proteomic alterations associated with clinical manifestation of ccRCC and define robust biomarkers for localized ccRCC and mRCC. Finally, we demonstrate the potential utility of these biomarkers for patient monitoring through proof-of-concept validation, establishing a foundation for liquid biopsy-driven prognostication and real-time ccRCC surveillance in clinics.

## Results

### ccRCC Plasma Cohort Study Design

Our study included plasma samples from 105 patients diagnosed with ccRCC between 2011 and 2022 (N = 105). Patients with confirmed ccRCC diagnosis and from whom sufficient sufficient plasma sample volume (1mL) was available pre-surgery (PRE-OP) and during follow-up (FU) time points after surgery (Figure 1A) were included, yielding 337 plasma samples.

**Figure 1.**
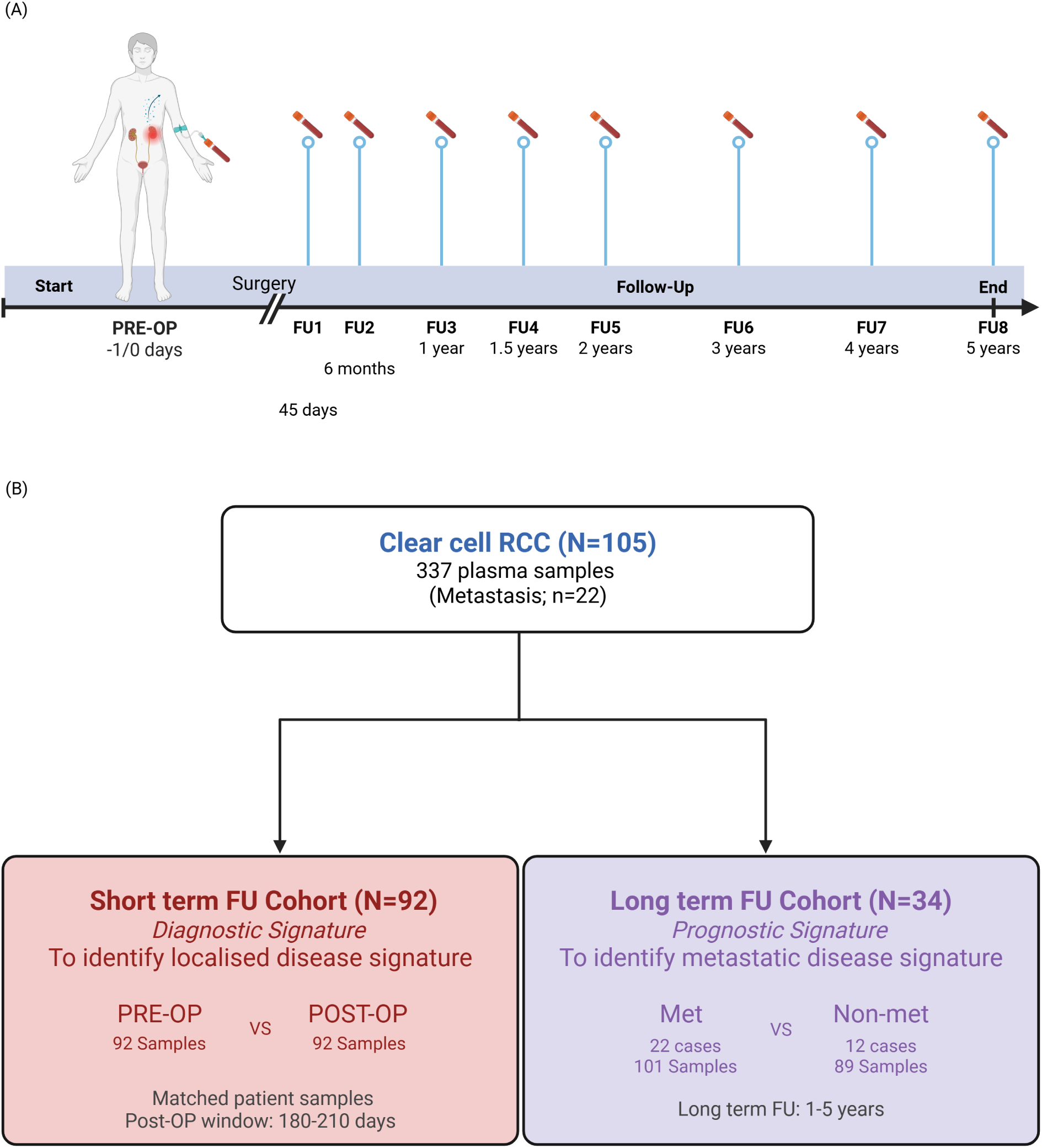
ccRCC liquid biopsy biomarker study design (A) Patient blood is collected before (PRE-OP) and at several follow-up (FU) time points after surgery. Processed plasma samples are used in this study. (B) Cohort selection and study design detailing the inclusion and exclusion criteria employed in ccRCC Cohort.

In this investigation, we further stratified our cohort into two distinct branches to address two primary research questions (Figure 1B). Short-term FU cohort (N = 92) was delineated to facilitate a comparative analysis of pre-surgery (PRE-OP) and post-surgery (POST-OP) plasma samples, aiming to identify diagnostic biomarkers for localized ccRCC. Therefore, this cohort included plasma samples from patients diagnosed with ccRCC localized to the kidney and who presented without known synchronous metastasis at diagnosis. In contrast, the Long-term FU cohort (N = 34) was established to examine longitudinal plasma samples to develop a signature indicative of disease dissemination or recurrence. This cohort comprised: (i) patients diagnosed with synchronous mRCC at first presentation in clinic, (ii) patients from the Short- term FU cohort who developed late metastasis during follow-up, and (iii) patients from the Short-term FU cohort with high-grade ccRCC without evidence of metastasis at the time of cohort selection. High-grade non-metastatic samples were used for comparison against mRCC samples since these advanced tumors often evolve into aggressive metastatic disease over time. Baseline characteristics of the patient population, including age, sex, tumor grade, and stage are comprehensively summarized in Table 1.

**Table 1.** Clinicopathological features of ccRCC patients included in the cohort.

| ccRCC Cohort (N= 105) |  |
| --- | --- |
| Sex (male/female) | 83/22 |
| Age, mean (SD) | 61.87 (14.36) |
| ISUP tumor grade |  |
| 1 | 9 |
| 2 | 51 |
| 3 | 27 |
| 4 | 18 |
| TNM stage |  |
| pT1a/b | 66 |
| pT2a/b | 7 |
| pT3 | 31 |
| pT4 | 1 |
| Metastasis (synchronous or late) | 22 |

ISUP, International Society of Urologic Pathologists

### Characterization of depleted ccRCC patient plasma

From the 337 ccRCC plasma samples, by immunoaffinity depletion of highly abundant proteins (see methods section) and employing mass spectrometry using data independent acquisition, we quantified 4024 unique proteins across all samples. On average, we quantified about 2400 proteins in each sample. The large cohort size necessitated the analysis of samples across eight batches, with observed batch effects subsequently corrected (Supplementary FigureS1A, C). The median biological variation between samples in our dataset ranged from 28.5% to 32.9%, attributable to anticipated inter-patient variability commonly observed in clinical settings, and was slightly higher than the median technical variation observed in quality control samples which was notably low, ranging from 9.4% to 10.6% (Supplementary FigureS1B). Importantly, unsupervised clustering of the dataset revealed no separation of samples based on clinical parameters such as stage, grade, sex or survival outcomes, highlighting an unbiased cohort design (Figure 2A). Further, analysis of variance confirmed that there was no remarkable separation of samples due to clinical parameters such as metastasis, despite the small variance observed in our dataset as expected in such large studies involving patient samples collected across different clinical time points (Supplementary FigureS1D-G). Notably, in our cohort, we achieved broad coverage of the dynamic range of the known plasma and whole human proteome as published in PaxDb 5.0 (Figure 2B, 2C), including canonical proteins and non-canonical isoforms. Our study quantified 24% and 11%, respectively, of the human plasma and whole human proteome (Figure 2D). This is comparatively lower than the average coverage observed in other tissue proteomic datasets deposited on PaxDb, potentially reflecting the inherent differences associated with plasma as a medium of investigation compared to the predominantly cellular proteins reported in public databases.

**Figure 2.**
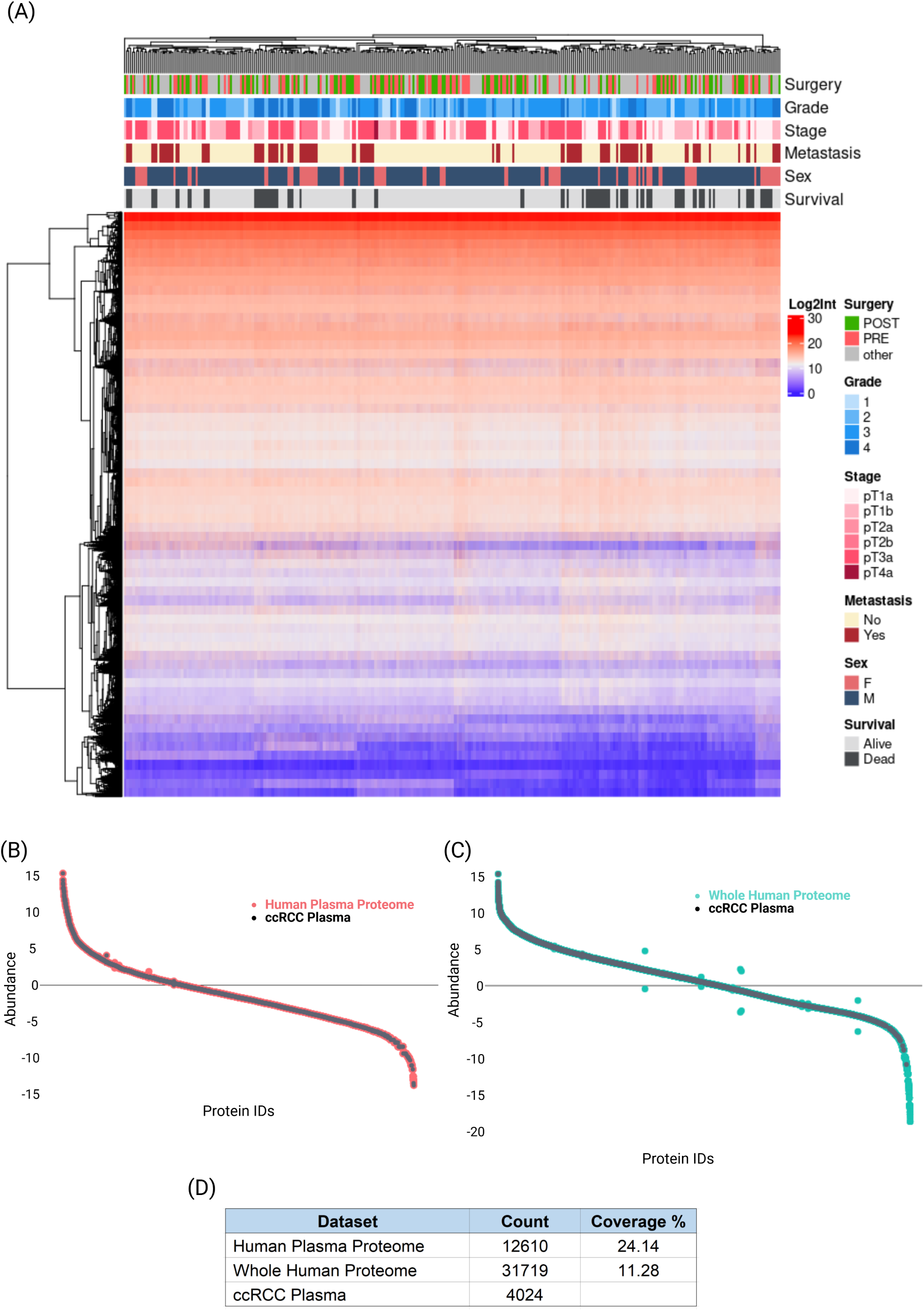
Exploratory analysis of ccRCC cohort patient plasma samples (A) Clustering of samples and heatmap of log transformed intensities of all proteins identified in ccRCC cohort. Rank plot of abundance of identified proteins in (B) human plasma proteome (orange trace) and (C) whole human proteome (blue trace). Black data points are proteins identified in the study. (D) Overview of identified protein count and coverage of dynamic range.

### Network analysis of ccRCC plasma reveals regulatory protein modules involved in disease biology and progression

To provide a comprehensive resource for the study of ccRCC molecular landscape in circulation, we first performed an analysis of the entire plasma proteome dataset. We constructed a weighted gene correlation network (WGCNA) of our cohort to examine and capture the protein expression and biological variability related to the observed clinical parameters corresponding to the ccRCC phenotype. We successfully identified 23 protein modules (Supplementary Figure S2C), which clustered based on similar differential co-expression. Module sizes ranged from 36 proteins in Module 11 (Dark Green) to 921 proteins in Module 23 (Grey) (Supplementary FigureS2B). Modules were correlated with relevant clinicopathological parameters to identify associations with localized disease, tumor progression, and metastasis and uncover biological implications of the circulating proteomic landscape. Several modules were identified with differing correlation strengths (Figure 3A). Specifically, Module 21 (Brown) was significantly associated with metastatic ccRCC patient samples (r = 0.32, p-value = 4e-09), and patient survival (r = 0.43, p-value = 4e-16), indicative of aggressive and advanced disease. Module 18 (Midnight Blue) was linked with metastatic ccRCC patient samples (r = 0.2, p-value = 3e-04) in addition to a strong association with the actual metastatic time points in the longitudinal study (r=0.19, p-value= 7e-04), serving as a potential indicator of early dissemination in high-risk patients. Additionally, Module 22 (Red) was associated with survival (r = 0.23, p-value = 3e-05) and Module 8 (Dark Red) was weakly associated with the presence of tumor before surgical resection (r = 0.15, p-value = 0.006).

**Figure 3.**
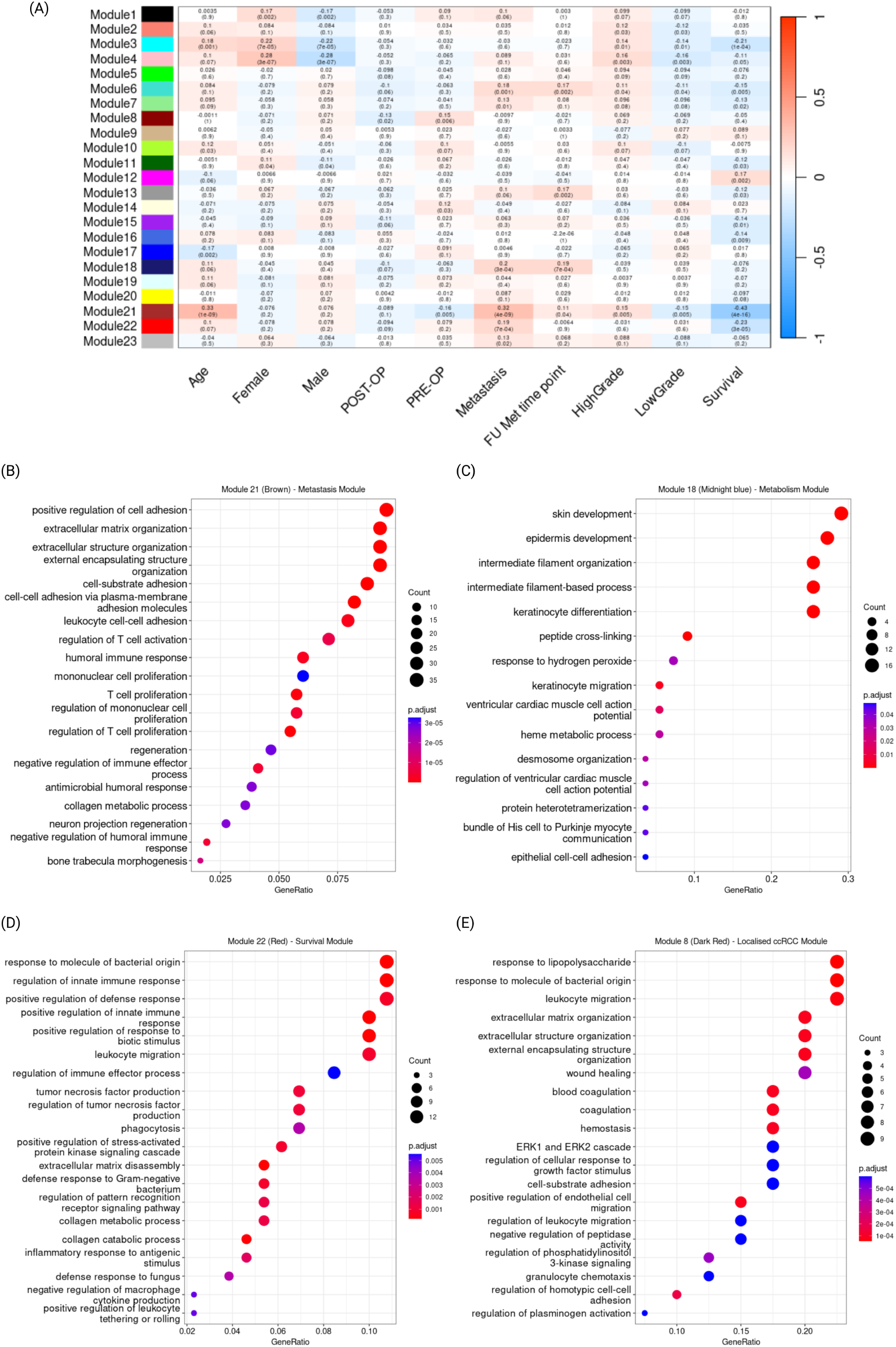
ccRCC plasma proteome network highlights key modules involved in ccRCC disease (A) Heatmap plot of Pearson’s correlations between network module eigengenes and key clinical parameters. Red indicates positive, and blue indicates negative correlation. Dot plots of Gene Ontology (GO) enrichment analysis showing top 20 significant biological processes identified in (D) Metastasis module (Module 21), (E) Metabolism module (Module 18), (F) Survival module (Module 22) and (G) Localized ccRCC module (Module 8). Color of dot represents the FDR of each term and size of dot represents gene count of the term involved. FDR, false discovery rate.

Functional enrichment analysis of the clinically relevant modules demonstrated the integral involvement of these proteins to the development of the ccRCC phenotype, corroborating findings from tumor tissue studies. Module 21 was strongly implicated in extracellular matrix reorganization-related processes, enabling disease dissemination and metastasis, while Module 18 largely contained proteins contributing to disrupted metabolism, specifically heme metabolism (Figure 3B-D). The metastasis Module 21 proteins included matrix metalloproteinase 2 (MMP2), collagen alpha 1 (COLA1), vascular cell adhesion molecule-1 (VCAM1), and homing cell adhesion molecule (HCAM/CD44), all known to play a central role in promoting the tumor microenvironment (TME) plasticity (Figure 3B). The metabolism Module 18 (Figure 3C) comprised proteins vital to tumor cell viability, importantly, ATP binding cassette subfamily B member (ABCB7), Heme oxygenase (HMOX), and Uroporphyrinogen III Synthase (UROS), [22, 25, 26]. These proteins are implicated in dysregulated heme metabolism or porphyrin overdrive, which further activates hypoxic signaling through the VHL-HIF pathway [27]. Further, survival Module 22 was defined by proteins involved in immune defense and inflammatory responses, accelerating cell death mechanisms (Figure 3D). Importantly, Module 8, correlating to localized ccRCC, contained proteins Thrombospondin 1 (THBS1), Angiopoietin 1 (ANGPT1), and Serglycin (SRGN), linked to processes such as cellular migration, and PI3K and ERK signaling pathways (Figure 3E), underscoring their value as potential biomarkers. Taken together, we identified distinct protein modules in the plasma proteome, underpinning ccRCC tumorigenesis and progression. Further mechanistic studies of these proteins could serve the identification of important molecular mechanisms in ccRCC.

### Differential expression of proteins in ccRCC patient plasma before surgery in Short-term FU Cohort

To explore alterations in the plasma proteome associated with ccRCC and identify biomarkers for localized disease, we compared plasma samples from ccRCC patients collected PRE-OP and POST-OP in the Short-term FU Cohort. Our study identified 50 proteins that exhibited differential expression following surgery (Supplementary Table S1).

Notably, 39 of these proteins were found to be significantly more abundant in PRE-OP samples, while a localized tumor was present (Figure 4A), potentially supporting tumor growth. Further examination through functional enrichment analysis highlighted the role of these 39 proteins in angiogenic and immune responses, apart from the involvement in dysregulated Wnt signaling, cellular metabolism and apoptotic cell death (Figure 4B). Importantly, we identified proteins such as Multimerin 1 (MMRN1), THBS1, Serum amyloid A1 (SAA1), Platelet factor 4 (PF4), Synaptotagmin (SYTL4), Glycoprotein Ib platelet subunit beta (GP1BB), Elastin microfibril interfacer 1 (EMILIN1), Envoplakin (EVPL), Myosin-9 (MYH9) and Myosin light chain 9 (MYL9) that are involved in processes linked to wound healing, now increasingly recognized to subvert similar molecular mechanisms as in cancer development and metastasis [28, 29]. Given the highly angiogenic characteristics of ccRCC, it is noteworthy that our study identified Prolactin (PRL), known for its pro-angiogenic roles in various cancers, and THBS1, which regulates angiogenesis in a context- and cell-specific manner, in the plasma of ccRCC patients in the Short-term FU Cohort.

**Figure 4.**
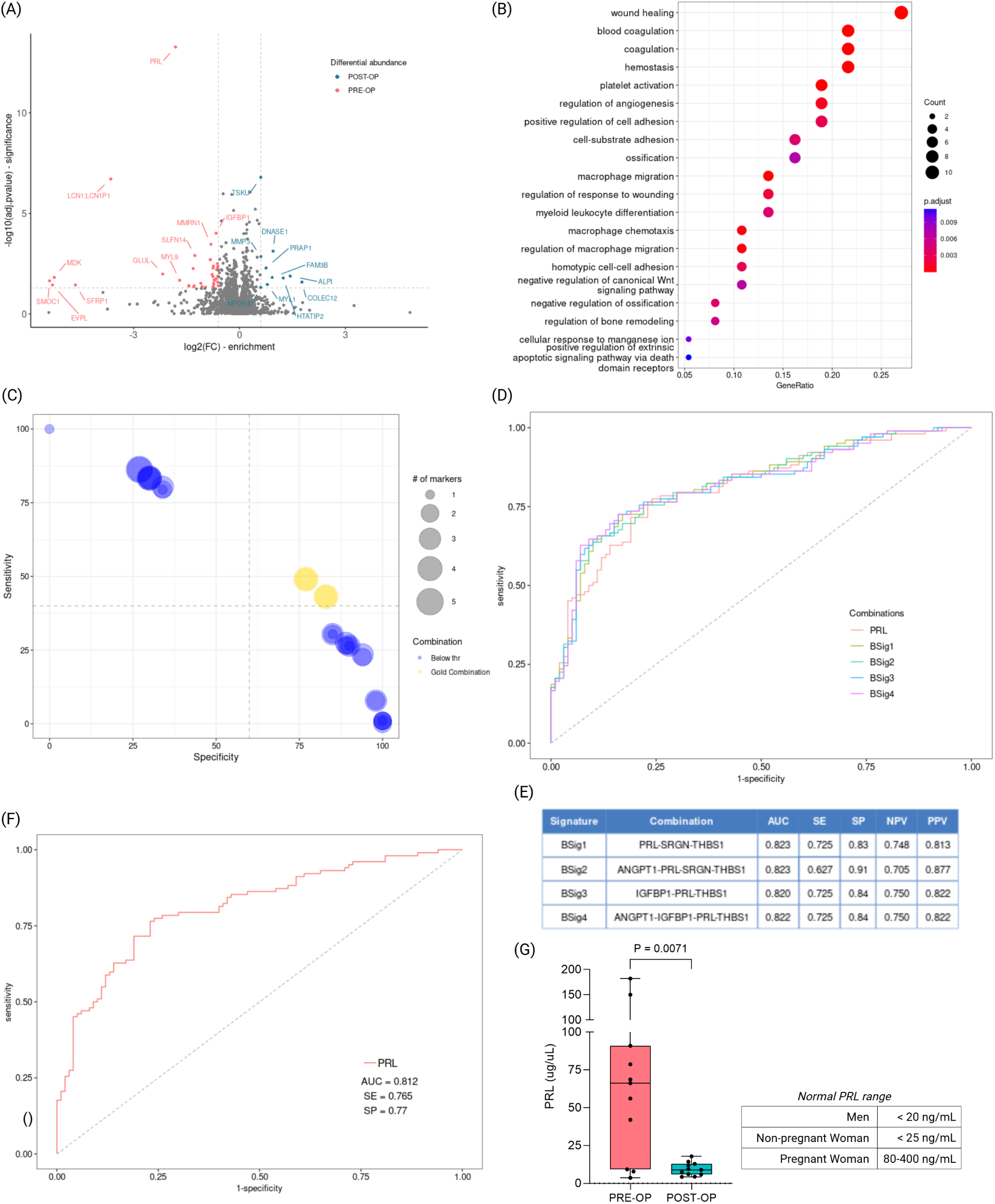
Differentially expressed proteins in PRE-OP ccRCC patient plasma samples play a role in critical oncogenic molecular mechanisms (A) Volcano plot of differentially expressed proteins, higher PRE-OP (orange) and POST-OP (blue) in Short-term FU Cohort. Significant hits were selected at a fold change cutoff of 1.5. (B) Dot plots of Gene Ontology (GO) enrichment analysis showing top 20 significant biological processes. Color of dot represents the FDR of each term and size of dot represents gene count of the term involved. (C) Bubble plot of sensitivity vs specificity of 26 combinations of five protein markers tested. Gold combinations include combinations of proteins crossing sensitivity (>=40) and specificity (>=60) threshold (D) ROC curve and (E) analysis metrics for selected protein signatures Bsig1, Bsig2, Bsig3, Bsig4 (F) ROC curve for PRL (G) Boxplot of PRL levels in a small validation cohort (N=11). FDR, false discovery rate; AUC, area under the curve; SE, sensitivity; SP, specificity; PPV, positive predictive value; NPV, negative predictive value

### Defining protein biomarkers and signatures reveals PRL as a promising biomarker for localized ccRCC

Among these 39 differentially expressed proteins, we prioritized the top 20 most significant proteins as potential biomarkers (Supplementary tableS1), with PRL being the most significantly expressed protein in PRE-OP plasma samples from patients with localized ccRCC in the Short-term FU cohort. Combining multiple proteins as biomarkers into a panel is a commonly employed strategy to improve diagnostic accuracy while simultaneously capturing a broader range of molecular mechanisms implicated in disease progression. Towards this end, we selected four proteins from the top 20 significant proteins with previously established roles in cancer, to include alongside PRL: THBS1 which is downregulated in ccRCC due to epigenetic silencing contributing to sunitinib resistance [30], Insulin-like growth factor-binding protein-1 (IGFBP1) [31], pro-angiogenic Angiopoietin (ANGPT1) [32] and Serglycin (SRGN) [33] that facilitate immune response in the TME. PRL, IGFBP1, THBS1, ANGPT1 and SRGN exhibited significant differences in abundance between PRE-OP and POST-OP plasma samples (Supplementary Figure S3).

A combinatorial receiver-operating characteristic curve (ROC) analysis of these five proteins allowed the identification of different permuted signatures for localized disease. We investigated 26 protein combinations from PRL, IGFBP1, THBS1, ANGPT1 and SRGN. We subsequently selected four combinations based on established thresholds for sensitivity (>= 40) and specificity (>= 60) (Figure 4C). The combination of PRL-SRGN-THBS1 (BSig1) had the best performance, with a high AUC (0.823) at 72.5% sensitivity, 83% specificity and a notable 74.8% positive predictive value (PPV) and 81.3% negative predictive value (NPV) (Figure4D). BSig 2 (ANGPT1- PRL-SRGN-THBS1) had a similar AUC (0.823) and higher specificity (91%) and PPV (87.7%), but lower sensitivity (62.7%) and NPV (70.5%). Signatures IGFBP1-PRL-THBS1 (BSig 3) and ANGPT1-IGFBP1-PRL-TBHS1 (BSig 4) had similar performance metrics with 72.5% sensitivity, 84% specificity, 82.2% PPV and 75% NPV. However, BSig 4 had a higher AUC (0.822) compared to BSig 3 (0.82) (Figure 4E). Interestingly, angiogenic proteins PRL and THBS1 were common in all the different signatures.

Since PRL was the most highly-differentially abundant protein in the Short-term FU Cohort, we assessed the utility of PRL as a liquid biopsy biomarker for ccRCC. PRL effectively discriminated plasma samples with localized tumors, achieving an excellent AUC (0.812) at 76.5% sensitivity (SE) and 77% specificity (SP) (Figure 4F). Furthermore, in order to substantiate PRL as a biomarker for ccRCC, we tested circulating plasma PRL levels in 11 patients from the Short-term FU cohort using a clinically approved immunoassay as an independent technical validation (patient clinicopathological characteristics detailed in Table 2). We observed significant changes in PRL levels correlating with localized tumors in PRE-OP samples, which exhibited up to a tenfold increase relative to POST-OP levels (p =0.0071), considerably surpassing the clinically defined ranges for healthy men (< 20 ng/mL) and healthy non-pregnant women (< 25 ng/mL) (Figure 4G).

**Table 2.** Clinicopathological characteristics of patients included for PRL validation.

| PRL Validation Cohort (N=11) |  |  |  |  |
| --- | --- | --- | --- | --- |
| Patient ID | WHO/ISUP Grade | Stage | Sex | Age |
| 130 | 3 | pT1a | f | 81 |
| 155 | 3 | pT3a | m | 49 |
| 158 | 2 | pT1a | f | 46 |
| 216 | 4 | pT3a | m | 53 |
| 221 | 2 | pT1b | f | 67 |
| 223 | 3 | pT3a | f | 69 |
| 249 | 2 | pT1a | f | 76 |
| 270 | 4 | pT3a | m | 55 |
| 291 | 4 | pT2b | f | 52 |
| 371 | 1 | pT1a | m | 47 |
| 391 | 2 | pT1a | m | 32 |

### Longitudinal differential expression of plasma proteins in metastatic patients in Long-term FU Cohort

Capturing the molecular landscape from longitudinally collected plasma samples facilitates the monitoring of early metastatic dissemination and recurrence of disease. Long-term FU cohort (N = 34, n= 190) was established to enable a comparison of (i) mRCC patients who exhibit synchronous metastasis at initial diagnosis or late metastasis during disease progression at subsequent follow-up, against (ii) ccRCC patients with high-grade tumors and no known history of metastasis at the time of study design. Using a rank product approach, we identified eight proteins with differential expression between the two groups: Complement component 1s (C1S), Complement component 2 (C2), F-Box and leucine rich repeat protein 17 (FBXL17), Transferrin receptor (TFRC), Insulin-like Ggowth factor binding protein 3 (IGFBP3), RAS guanyl releasing protein 2 (RASGRP2), Alpha-1-microglobulin/Bikunin precursor (AMBP), Ficolin 3 (FCN3) (Figure 5A). For further biomarker exploration, six proteins, C1S, C2, TFRC, IGFBP3, RASGRP2, and AMBP, were chosen due to their significantly increased expression in metastatic samples compared to high-grade ccRCC plasma samples (Figure 5B). Notably, these six biomarker candidates exhibited higher expression in mRCC patients with late metastasis than in the high-grade, non-metastatic ccRCC group, with a change in their expression around the clinical detection of metastasis (Figure 5C).

**Figure 5.**
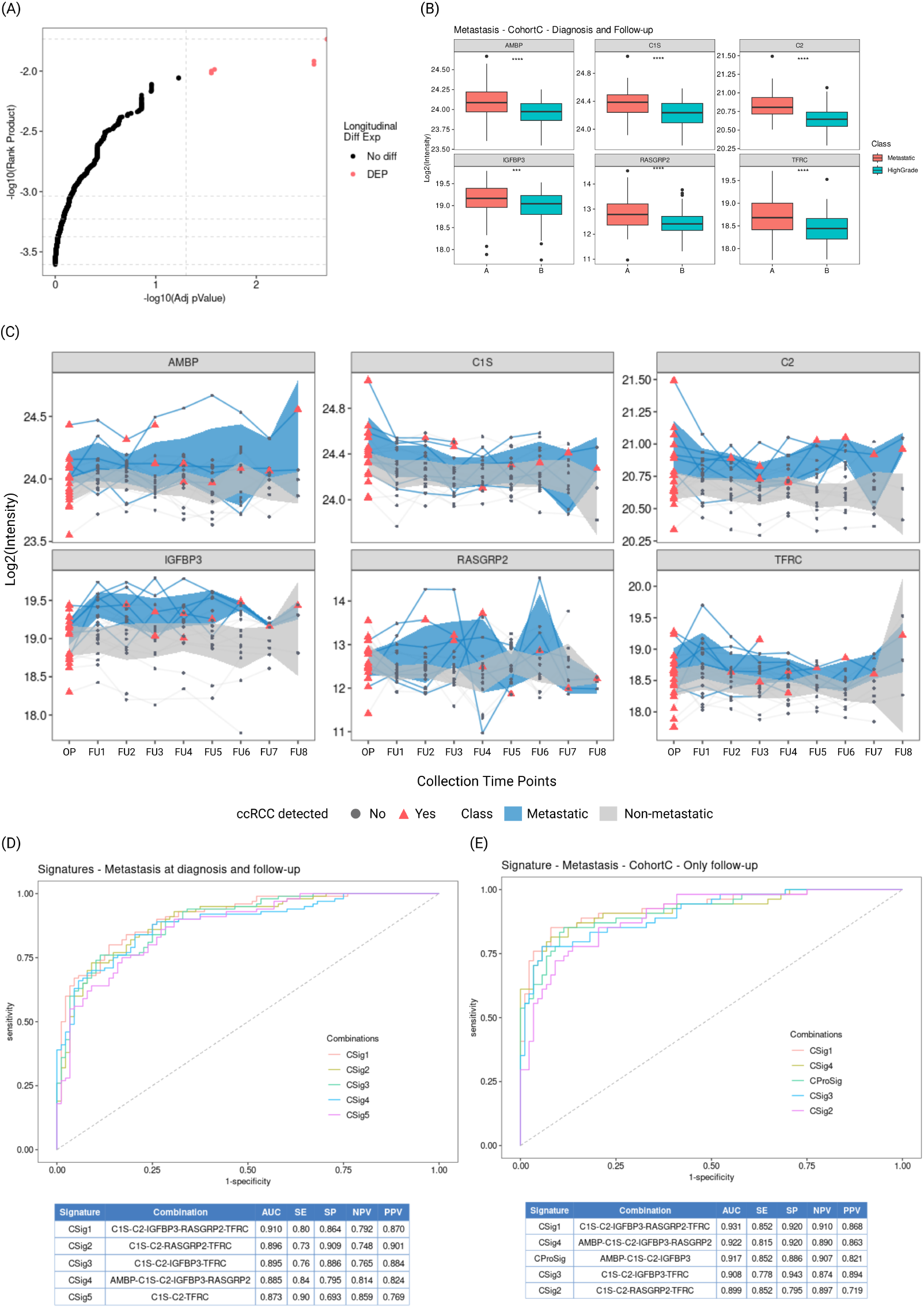
Longitudinal analysis of metastatic samples reveals six differentially expressed proteins and their combinations as potential prognostic signatures in Long-term FU Cohort (A) Scatter plot of rank product vs adjusted p-value. Proteins with the highest rank product and adj p-value < 0.05 are highlighted (B) Boxplot showing distribution of six markers - AMBP, C1S, C2, IGFB3, RASGRP2, and TFRC, across longitudinal samples from metastatic (orange) and non-metastatic, high-grade (blue) samples (C) Expression of six markers across follow-up time points in metastatic (blue) and non-metastatic (grey) ccRCC patients. Line indicates expression of proteins in individual patients. Ribbon indicates 95% confidence interval of expression. Time points when ccRCC is detected (PRE-OP or during follow-up, FU) are highlighted in orange triangles (D) ROC curve and analysis metrics for selected protein signatures CSig1, CSig2, CSig3, CSig4, CSig5 modelled comparing all metastatic patients vs high-grade (E) ROC curve and analysis metrics for selected protein signatures CSig1, CSig4, CProSig, CSig3, CSig4 modelled in patients with metastasis at prognosis. ROC DEP, differentially expressed protein; AUC, Area under the curve; SE, sensitivity; SP, specificity; PPV, positive predictive value; NPV, negative predictive value

### Defining protein biomarkers and signatures for metastatic ccRCC

We assessed combinations of these markers to evaluate their significance as metastatic signatures for all mRCC patients, including both synchronous and late metastasis (CSig, Figure 5D). A total of 63 combinations from these six candidate proteins were analysed, yielding 16 that achieved the threshold for sensitivity (>= 40) and specificity (>= 60). The top five combinations with the highest AUC were selected as potential signatures. CSig1 (C1S-C2- IGFBP3-RASGRP2-TFRC) demonstrated the highest performance, achieving an AUC (0.91) at 80% sensitivity and 86.4% specificity and a clinical PPV of 87% (Figure 5D). CSig2 (C1S- C2-IGFBP3-RASGRP2-TFRC) and CSig3 (C1S-C2-IGFBP3-TFRC) were comparably effective, with AUC of 0.896 and 0.895, respectively, although CSig2 had a higher PPV of 90.1%. CSig4 (MBP-C1S-C2-IGFBP3-RASGRP2) recorded an AUC of 0.885 at 84% sensitivity and 79.5% specificity, while CSig5 (C1S-C2-TFRC) performed with the lowest AUC of 0.873.

We then tested the prognostic capability of the six proteins to identify ccRCC patients developing only late metastasis. Intuitively, the top five performing combinations in this analysis included CSig1, CSig2, CSig3 and CSig4 observed in mRCC patients with synchronous metastasis at clinic (Figure 5E). CSig1 remained the best performing prognostic signature, showing improved performance with an AUC of 0.931 and increased sensitivity (85.2%) and specificity (92%) in detecting late metastasis. Additionally, CProSig (AMBP-C1S-C2-IGFBP3) emerged as a unique signature for late mRCC samples, achieving an AUC of 0.917 at a clinical PPV of 82.1%. Interestingly, C1S and C2 were consistently included in the defined CSig combinations, demonstrating a strong correlation with disease progression in both synchronous and late mRCC patients.

## Discussion

The identification of effective circulating protein biomarkers in ccRCC patient plasma is fraught with challenges arising from the broad dynamic range of plasma proteins, where a limited number of highly abundant proteins obscure the detection of lower-abundance proteins potentially linked to disease pathology. Although some studies have explored circulating tumor DNA (ctDNA) as a biomarker, ccRCC is notably a ctDNA-low malignancy, with standard sequencing methodologies detecting ctDNA in only about 27.5% of patients [34]. This limitation underscores the difficulty of translating ctDNA-based diagnostics into routine clinical practice and highlights the need for complementary protein biomarker strategies. The challenge is compounded by the scarcity of large, representative longitudinal patient cohorts, which are essential for discovering markers that can predict late metastasis during the course of ccRCC progression [11].

To address these gaps, our study analyzed an extensive longitudinal ccRCC plasma cohort, presenting one of the most comprehensive proteomic datasets currently available for this disease. Unlike previous studies that compared PRE-OP ccRCC samples with healthy individuals, our approach focused on matched samples before and after tumor removal, and included extended longitudinal follow-up samples [35, 36]. Plasma samples collected between 180 and 210 days post-surgery were included as POST-OP samples, ensuring clearance of transient surgical signatures and allowing a clearer snapshot of ccRCC dynamics and progression. This design is critical, as surgical intervention can induce significant changes in the blood proteome, notably increasing the levels of angiogenic proteins and enhancing wound-healing processes in various cancers for up to four weeks post-surgery [37]. Additionally, we employed depletion strategies to remove highly abundant plasma proteins, enabling the detection and characterization of disease-relevant, lower-abundance proteins that would otherwise be missed by conventional native plasma proteomics [19]. This refined approach resulted in increased protein identifications and enabled a robust network-based analysis to elucidate the role of the circulating proteome in both the initiation and dissemination of ccRCC.

Our network analysis provides a comprehensive atlas of the ccRCC plasma proteome and contextualizes the subsequent identification of biomarker candidates. We identified four distinct modules (Module 21, 18, 22, and 8) significantly enriched for proteins involved in the metastatic cascade and the metabolic reprogramming, characteristic of ccRCC. Although various malignancies share perturbations foundational to oncogenesis, the clinically relevant modules identified in our study reflect a ccRCC-specific molecular landscape, perpetuated by the hallmark dsyregulated VHL-HIF axis and lipid metabolism [38], with several therapeutic strategies currently being developed using these proteins. Disrupted glycolysis, lipidogenesis, and glutamine synthesis are established hallmarks of ccRCC, while heme metabolism is becoming increasingly recognized as significant for disease progress. Notably, HMOX1, an integral player in the heme metabolic pathways identified in our metabolism module, has recently been proposed as a tissue biomarker for predicting response to immunotherapy plus tyrosine kinase treatment in advanced RCC [39]. Furthermore, MMP2, VCAM1, CD44 and COL1A1 identified in the metastatic module 21, are well-known for their roles in cancer metastasis and have been explored as therapeutic targets across multiple solid tumors [40–43]. Specifically, MMP2, VCAM1, and CD44 are associated with poorer patient outcomes, mediating proliferation, migration, and immune responses through ERK/MAPK [44], VHL-HIF [45] and NF*κ*B [46] signaling pathways respectively. Although the precise molecular mechanisms in ccRCC require further establishment, *COLA1* is known to be hypomethylated in tumor tissue, enabling its transcriptional over-expression, possibly contributing to the elevated circulating COLA1 observed in plasma [47]. Collectively, we provide evidence that the ccRCC-specific molecular changes observed at tissue level are mirrored in patient blood, thereby presenting an extensive and clinically relevant map of the plasma proteomic landscape for advancing liquid biopsy biomarker discovery and further mechanistic investigations in ccRCC.

Our study identified PRL, THBS1, ANGPT1, IGFBP1, and SRGN as promising plasma biomarkers for distinguishing localized ccRCC. Interestingly, decreased THBS1 and increased SAA1 expression in ccRCC tumor tissue predict more aggressive and advanced disease [30, 48], with SAA1 promoting pro-restitutive epithelial phenotype and THBS1 promoting cell migration through matrix remodeling, mechanisms relevant to both wound healing and tumor progression [49, 50], underscoring their utility as biomarkers. Among these, PRL stood out as an effective single biomarker for localized ccRCC, consistent with its known role as a peptide hormone crucial for mammary gland development and lactation, and promoting proliferation, angiogenesis, migration, and invasion in hormone-dependent cancers [51]. ccRCC is increasingly recognized as a hormonally active malignancy with higher incidence in males, and despite the sexually dimorphic expression and the intricate involvement of PRL in female health, our data revealed no significant sex-based differences in plasma PRL, supporting its broad applicability as a diagnostic marker (Supplementary FigureS3C). While Yu et al. correlated Follicle-Stimulating Hormone (FSH) and Estradiol to ccRCC progression in males, they did not detect elevated PRL, likely due to unclear sampling times in the study design [52]. In contrast, our results demonstrated a marked increase in PRL before surgery in both male and female patients with localized tumors, returning to normal levels postoperatively, reinforcing PRL as an effective biomarker to include during initial diagnosis.

A 1981 study reported a modest preoperative increase in serum PRL in ccRCC patients, although not significantly linked to pathology [53], providing early but unexplored evidence for PRL in ccRCC biology. More recently, Yang et al. highlighted an increase in PRL in ccRCC tissue, suggesting it facilitates tumor fibrosis and progression via the JAK/STAT signaling pathway [54]. Furthermore, PRL is known to promote proliferation, migration, and invasion in various cancers by activating oncogenic AKT, MAPK, and JAK/STAT signaling, compelling the development of therapeutic strategies targeting the PRL receptor [51]. Our findings recommends the importance of recognizing hormonal and endocrinal abnormalities as important and indicative biomarkers for ccRCC, rather than considering paraneoplastic syndromes as miscellaneous for clinical diagnosis. In ccRCC, an interplay between PRL and VHL-HIF axis could contribute to the adaptive metabolic changes in the tumor microenvironment, though further studies are necessary to clarify the exact molecular mechanisms. Additionally, key biomarker for localized ccRCC, PRL, THBS1, and IGFBP1 exhibit interdependent expression in other tumor types and may influence stromal differentiation, meriting deeper investigation of such cooperative mechanisms in ccRCC biology [55].

We also identified AMBP, C1S, C2, IGFB3, RASGRP2, and TFRC as biomarker candidates for both synchronous and late mRCC. Notably, these six proteins appear especially promising when we consider their potential as prognostic biomarkers for the detection of late metastasis and minimal residual disease, as shown by the distinct change in their plasma abundance precisely at the time metastasis is detected in clinic. In our retrospective longitudinal cohort, their differential expression clearly distinguishes patients developing late metastasis from those with aggressive high-grade ccRCC who did not progress to develop metastasis, underscoring their strong prognostic potential and clinical utility. All effective signature combinations in patient plasma included C1S and C2, highlighting their potential importance. While previous research showed elevated levels of C1S and C2 in ccRCC patients compared to healthy controls without prognostic significance [56], our findings provide novel evidence suggesting these proteins may be valuable for detecting early metastatic spread and minimal residual disease, pending rigorous validation using independent immunoassay. The role of complement signaling in modulating immunosuppressive metastatic niche and supporting organotropism, as demonstrated in breast cancer patients receiving chemotherapeutic intervention [57], remains unexplored in ccRCC and warrants further investigation to determine the extent of involvement of complement signaling in late metastasis, recurrence and minimal residual disease. While we did not identify any overlap between the biomarker candidates for localized and metastatic disease, the presence of IGFBP family proteins, namely IGFBP1 in localized signatures and IGFBP3 in metastatic signatures, suggests a shared molecular underpinning. IGFBP1 and IGFBP3 are both located on chromosome 7, and this increased expression observed in plasma could correspond to a functional outcome of the frequently observed chromosome 7 trisomy in ccRCC [58]. Additionally, THBS1 functions as an anti-angiogenic protein activated by IGFBP3 in ovarian cancer, although such a regulatory axis remains to be defined in ccRCC [59].

Taken together, our findings underscore the functional relevance of plasma proteins as minimally invasive biomarkers for both diagnosis of localized ccRCC and the prognosis of synchronous and late metastasis. However, as with all biomarker discovery efforts, rigorous validation in larger, independent cohorts, and translation from high-resolution proteomics to easily implementable immunoassays. Our proof-of-concept validation of PRL using clinically approved immunoassays provides critical first evidence for the easy implementation of these biomarkers in routine practice. By defining biomarker signatures associated with ccRCC pathology, we demonstrate the potential of liquid biopsy as a minimally invasive technology for the improved longitudinal management of ccRCC patients, especially those at high risk for metastasis. This work lays a strong foundation for integrating proteomic biomarkers into routine clinical practice, ultimately advancing the care and outcomes for ccRCC patients.

## Methods

### USZ Plasma Sample Collection

The Cantonal Ethics Committee, Zürich, Switzerland, approved this plasma collection and study protocol (BASEC 2019-01959). The Biobank at the Department of Pathology and Molecular Pathology, University Hospital of Zurich provided the plasma samples. All patients provided written consent for this study. Plasma samples were collected from patients diagnosed with ccRCC before surgery and up to eight-time points after surgery, as shown in Figure 1A. Blood samples were collected in EDTA vacutainers and stored as plasma per a standardised sample processing protocol. After centrifugation at 3600*g* at 4°C for 15min, the samples were aliquoted in 1.5mL eppendorf tubes and stored at -80°C until further use.

### Sample Preparation

This study was conducted with experimental support from Biognosys AG. Samples were shipped frozen on dry ice to Biognosys AG. Plasma samples were depleted using High Select Top14 Abundant Protein Depletion Resin (Thermo Scientific) and further prepared on a Hamilton Microlab STAR Liquid Handling System according to Biognosys’ standardised protocol. Samples were reduced, alkylated and digested to peptides at 37°C using Trypsin (Promega, 1:100 protease to total protein ratio) and Lys C (Fujifilm Wako Chemicals, 1:200 protease to total protein ratio). Peptides were desalted using an HLB µElution plate (Waters Corporation) and dried down. Peptides were then resuspended with 1% acetonitrile/0.1% formic acid in water and spiked with Biognosys’ iRT kit calibration peptides. Peptide concentrations were determined with a Micro BCA assay (Pierce, Thermo Fisher Scientific).

### Hyper Reaction Monitoring (HRM) Mass Spectrometry Acquisition

For Data Independent Acquisition (DIA) LC-MS/MS measurements, 2 µg of sample peptides were injected on an in-house packed reversed phase column on a Thermo Scientific NeoVanquish UHPLC nano liquid chromatography system connected to a Thermo Scientific Orbitrap^TM^ Exploris 480^TM^ mass spectrometer equipped with a Nanospray Flex^TM^ ion source and a FAIMS Pro^TM^ ion mobility device (Thermo Scientific). LC solvents were (A) water with 0.1 % FA, (B) 80% acetonitrile, 0.1% FA in water. The non-linear LC gradient was 1- 50% solvent B for 172 minutes, followed by a column washing step in 90% B for 5 minutes, and a final equilibration step of 1% B for 1 column volume with a flow rate set to a ramp between 500 to 250 nL/min (min 0: 500 nL/min, min 172: 250 nL/min, washing at 500 nL/min). The FAIMS DIA method consisted per applied compensation voltage of one full range MS1 scan and 34 DIA segments as adopted from Bruderer et al. [60] & Tognetti et al.[24].

### Database search and quantitative analysis of HRM LC-MS/MS data

For whole proteome analysis, DIA mass spectrometric data were processed using the Spectronaut software (v18.1, Biognosys AG). The analysis was performed using default settings, including a 1% false discovery rate control at both peptide and protein levels and allowing for up to two missed cleavages. Carbamidomethyl was specified as fixed modification and N-term acetylation, methionine oxidation, ammonia loss, and deamidation (NQ) as variable modifications. Protein identifications were conducted using a UniProt (RRID:SCR_002380) .fasta database (*Homo sapiens*, 2023-07-01). For the library generation, the default Spectronaut settings were used, with deamidated peptides excluded at the library level. Data filtering was performed using row-based extraction. Normalisation of HRM measurements was carried out using local normalisation based on retention time (RT)-dependent local regression model as described by Callister et al.[61]. The sparse dataset consisted of precursors filtered based on q-value, with missing values estimated using background signal imputation. Any observed batch effects were corrected using the Hermes pipeline from Biognosys AG.

### Data Analysis

Distance in heat maps was calculated using the "Manhattan” method, and the clustering using "ward.D” for both axes. Principal component analysis (PCA) and partial least squares discriminant analysis (PLS-DA) was conducted in R using ropls (v1.36.0, RRID:SCR_016888), with standard scaling using the nipals algorithm. To test differential protein abundance in the Short-term FU Cohort, fold changes for each protein were analysed using a two-sample Student’s t-test. The following thresholds were applied for candidate identification: p value < 0.05; absolute average log2 ratio > 0.58 (fold change > 1.5). RolDE (v1.8.0), a composite method combining three differential analysis algorithms to derive a rank product, was used to calculate the longitudinal differential expression in the Long-term FU Cohort. The "Mixed1" model was selected for the analysis of non-aligned metastatic time points, allowing us to develop models with random effects for each individual. Estimated significance values were corrected for multiple testing using the Benjamini-Hochberg method, and candidates with adjusted p-value < 0.05 were selected for further analysis. Protein-protein interaction networks were constructed using String db (v12.0, RRID:SCR_005223, https://string-db.org) with a minimum interaction score of 0.04 (medium confidence). Gene Ontology (GO) enrichment analysis was performed in R using ClusterProfiler (v4.7.1; RRID:SCR_016884) using a human genome annotation file obtained from Org.Hs.eg.db (v3.17.0, RRID:SCR_024739). Figures were generally plotted in R using ggplot2 (v3.4.2; RRID:SCR_014601).

### Weighted Gene Co-expression Network Analysis (WGCNA)

We constructed a weighted gene co-expression network (RRID:SCR_003302) to identify and establish an association between ccRCC plasma proteins and derive a comprehensive overview of the different networks of proteins reflecting the ccRCC disease state in circulation (Supplementary FigureS2). The input for network construction was the spectral data matrix derived after significance value estimation, as described in the data analysis section above. An optimal soft threshold power *β* = 5 was used to calculate an adjacency and topological overlap matrix. Oncogenesis, disease progression and metastasis are governed by several mutually antagonising and complex regulatory molecular mechanisms, such as the compounded effect of the loss of tumor suppressor and gain of oncogene. Using an unsigned network approach allowed us to develop modules that were either positively or negatively correlated with relevant clinical parameters. Modules were constructed to automatically merge highly correlated proteins (deep split = 2, minimum module size = 30, cut height = 0.3), with each module assigned a distinct Color. The calculated eigengene summarizes each module’s primary expression pattern among its constituent proteins. To determine the association of modules with clinical traits, we calculated the Pearson correlation between the module eigengene and the relevant clinical parameters. We next calculated the gene significance (GS) to assess metastasis associations for selecting prognostic biomarker combinations. The gene data from these relevant modules were subsequently extracted for further analysis using CombiROC.

### Biomarker combinatorial analysis

We analysed multiple potential biomarkers to identify the best marker combinations using CombiROC (v0.3.4). For the combinatorial ROC analysis of each cohort (Short-term FU Cohort for localized ccRCC, Long-term FU Cohort for metastatic ccRCC), different proteins were selected, and a signal threshold was computed for the analysis respectively. Combinations with the highest AUC, sensitivity >= 40 and specificity >= 60, were identified as golden combinations for biomarker signatures.

### PRL Immunoassay

We validated PRL in eight ccRCC patients (individual patient characteristics listed in Table 2). This was performed with experimental support from the Institute of Clinical Chemistry, USZ. Briefly, frozen plasma samples were thawed, and a minimum volume of 100*µ*L was quantified using a fully automated cobas immunology analyser workflow using the approved *in vitro* diagnostic Elecsys® Prolactin II kit (F. Hoffmann-La Roche Ltd).

## Acknowledgments

The authors would like to thank all members of the Tissue Biobank, Department of Pathology and Molecular Pathology and the staff from the Department of Urology, University Hospital of Zurich for aiding the collection, processing and banking of patient blood samples. We are grateful to Biognosys AG for their support in proteomic data acquisition and preliminary analysis. We would also like to thank Katharina Spanaus from the Institute of Clinical Chemistry, University Hospital of Zurich for the support with validation experiments. This work was funded by the Swiss National Science Foundation (SNF-Project 31003_135792) and the Lotte und Adolf Hotz-Sprenger Stiftung.

