## Supplementary material for "Decoding the Plasma Proteomic Landscape of Clear Cell Renal Cell Carcinoma Reveals Diagnostic and Prognostic Liquid Biopsy Biomarkers": Supplemtary file

**Table S1.** Differentially expressed proteins in PRE-OP compared to POST-OP plasma samples in Short-term FU Cohort

| ID | Name | Description | log(FC) | DE | p-value |
|---|---|---|---|---|---|
| P01236 | PRL | Prolactin | -1.8142 | UP | 5.30E-14 |
| P31025 | LCN1 | Lipocalin-1;Putative lipocalin 1-like protein 1 | -3.6453 | UP | 1.95E-07 |
| P08833 | IGFBP1 | Insulin-like growth factor-binding protein 1 | -0.6593 | UP | 9.71E-05 |
| Q13201 | MMRN1 | Multimerin-1 | -0.8073 | UP | 3.44E-04 |
| P0C7P3 | SLFN14 | Protein SLFN14 | -1.2674 | UP | 1.23E-03 |
| P07996 | THBS1 | Thrombospondin-1 | -0.8239 | UP | 2.00E-03 |
| P09210 | GSTA2 | Glutathione S-transferase A2 | -0.6233 | UP | 3.14E-03 |
| Q15389 | ANGPT1 | Angiopoietin-1 | -0.7393 | UP | 4.16E-03 |
| P05067 | APP | Amyloid-beta precursor protein | -0.6106 | UP | 4.23E-03 |
| Q6UXH1 | CRELD2 | Protein disulfide isomerase CRELD2 | -0.6083 | UP | 4.78E-03 |
| P60660 | MYL6 | Myosin light polypeptide 6 | -0.7513 | UP | 5.25E-03 |
| P10451 | SPP1 | Osteopontin | -1.3099 | UP | 5.49E-03 |
| Q14766 | LTBP1 | Latent-transforming growth factor beta-binding protein 1 | -0.6406 | UP | 6.27E-03 |
| P35579 | MYH9 | Myosin-9 | -0.5997 | UP | 6.60E-03 |
| P15104 | GLUL | Glutamine synthetase | -2.1731 | UP | 1.04E-02 |
| Q96C24 | SYTL4 | Synaptotagmin-like protein 4 | -0.7762 | UP | 1.11E-02 |
| P10124 | SRGN | Serglycin | -0.6161 | UP | 1.34E-02 |
| P13224 | GP1BB | Platelet glycoprotein Ib beta chain | -0.7539 | UP | 1.38E-02 |
| Q9Y6C2 | EMILIN1 | EMILIN-1 | -0.5881 | UP | 1.44E-02 |
| P21741 | MDK | Midkine | -5.2432 | UP | 1.51E-02 |
| P21675 | TAF1 | Transcription initiation factor TFIID subunit 1 | -0.6696 | UP | 1.84E-02 |
| P63167 | DYNLL1 | Dynein light chain 1, cytoplasmic | -0.7092 | UP | 1.96E-02 |
| P24844 | MYL9 | Myosin regulatory light polypeptide 9 | -1.6948 | UP | 2.13E-02 |
| Q92747 | ARPC1A | Actin-related protein 2/3 complex subunit 1A | -0.7146 | UP | 2.19E-02 |

| | | | | | |
|---|---|---|---|---|---|
| Q9H4F8 | SMOC1 | SPARC-related modular calcium-binding protein 1 | -5.3824 | UP | 2.23E-02 |
| P02776 | PF4 | Platelet factor 4 | -0.6186 | UP | 2.66E-02 |
| Q00765 | REEP5 | Receptor expression-enhancing protein 5 | -1.0928 | UP | 3.01E-02 |
| P09603 | CSF1 | Macrophage colony-stimulating factor 1 | -0.7307 | UP | 3.08E-02 |
| Q92817 | EVPL | Envoplakin | -5.2983 | UP | 3.62E-02 |
| Q8N474 | SFRP1 | Secreted frizzled-related protein 1 | -4.6432 | UP | 3.67E-02 |
| O75369 | FLNB | Filamin-B | -0.7354 | UP | 3.78E-02 |
| Q9P0Z9 | PIPOX | Peroxisomal sarcosine oxidase | -1.4396 | UP | 3.84E-02 |
| P09496 | CLTA | Clathrin light chain A | -0.6057 | UP | 3.94E-02 |
| P0DJI8 | SAA1 | Serum amyloid A-1 protein | -1.2865 | UP | 4.01E-02 |
| P46952 | HAAO | 3-hydroxyanthranilate 3,4-dioxygenase | -1.4311 | UP | 4.04E-02 |
| P00325 | ADH1B | All-trans-retinol dehydrogenase [NAD(+)] ADH1B | -0.7442 | UP | 4.39E-02 |
| P10412 | H1-4 | Histone H1.4 | -1.0596 | UP | 4.39E-02 |
| Q92765 | FRZB | Secreted frizzled-related protein 3 | -1.2974 | UP | 4.44E-02 |
| Q14240 | EIF4A2 | Eukaryotic initiation factor 4A-II | -0.9619 | UP | 4.67E-02 |
| Q8WUA8 | TSKU | Tsukushi | 0.6066 | DOWN | 1.60E-07 |
| P24855 | DNASE1 | Deoxyribonuclease-1 | 0.9534 | DOWN | 7.53E-04 |
| P08254 | MMP3 | Stromelysin-1 | 0.6076 | DOWN | 1.39E-03 |
| Q96NZ9 | PRAP1 | Proline-rich acidic protein 1 | 0.7500 | DOWN | 5.17E-03 |
| P10645 | CHGA | Chromogranin-A | 0.5811 | DOWN | 1.14E-02 |
| P09923 | ALPI | Intestinal-type alkaline phosphatase | 1.4344 | DOWN | 1.30E-02 |
| P58499 | FAM3B | Protein FAM3B | 0.9267 | DOWN | 1.54E-02 |
| Q9BUP3 | HTATIP2 | Oxidoreductase HTATIP2 | 1.2407 | DOWN | 1.64E-02 |
| Q5KU26 | COLEC12 | Collectin-12 | 1.7677 | DOWN | 2.61E-02 |
| P05976 | MYL1 | Myosin light chain 1/3, skeletal muscle isoform | 0.7909 | DOWN | 3.45E-02 |
| Q08431 | MFGE8 | Lactadherin | 0.6003 | DOWN | 4.85E-02 |

FC, fold change; DE, differential expression;

UP, higher expression in PRE-OP samples; DOWN, lower expression in PRE-OP samples

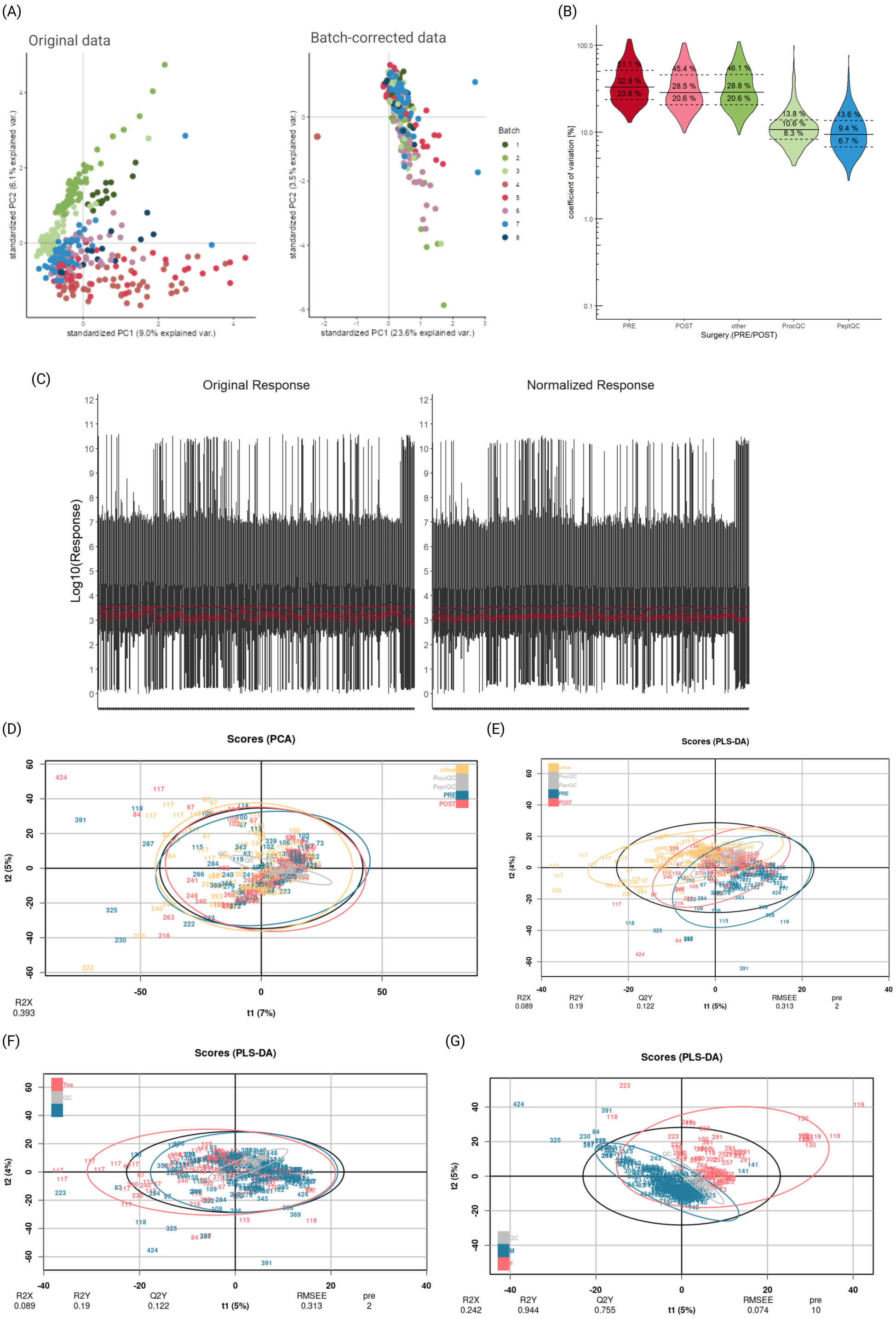


**Figure S1.** Data processing and quality control of depleted ccRCC patient plasma (A) PCA plots of variance in dataset before (left) and after (right) batch correction (B) Violin plot showing distribution of CV for batch-corrected protein quantifications grouped by time point of collection: PRE-OP (red), POST-OP (pink), follow-up (green), experimental controls (light green & blue) (C) Boxplot of log transformed protein abundance before (right) and after (left) normalisation. PCA (D) and PLS-DA (E) biplot of variance in dataset after batch correction and normalisation among PRE-OP (pink), POST-OP (blue), follow-up (yellow) and experimental control (grey) samples. PLS-DA biplot of variance in dataset grouped by metastasis (F) (metastasis - pink, local - blue, experimental controls - grey) and sex (G) (male - blue, female - pink, experimental control - grey). PCA, principal component analysis; PLS-DA, partial least squares discriminant analysis; CV, coefficient of variation.

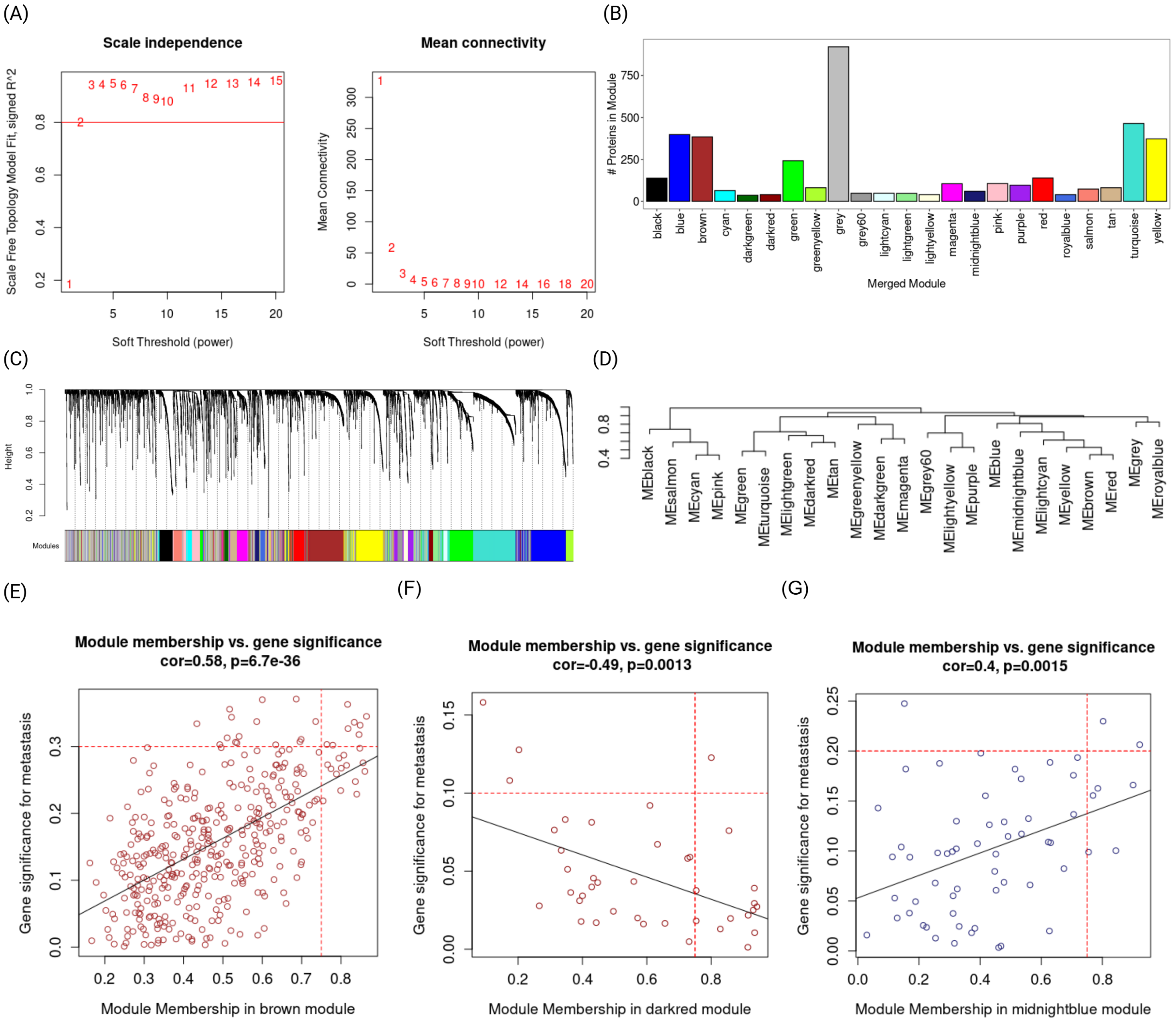


**Figure S2.** Network analysis of longitudinal ccRCC plasma (A) Scale-free topology fit index (left) and mean connectivity versus soft threshold power($\beta$). $\beta$= 5 selected for analysis (B) Barplot of proteins in each defined module (C) Protein clustering dendrogram based on TOM-based dissimilarity (D) Dendrogram of module eigengene clustering. Correlation between gene significance for metastasis and module membership for proteins in (E) Module 21 (Brown), (F) Module 18 (Midnight blue) and (G) Module 8 (Dark Red). TOM, topological overlap matrix.

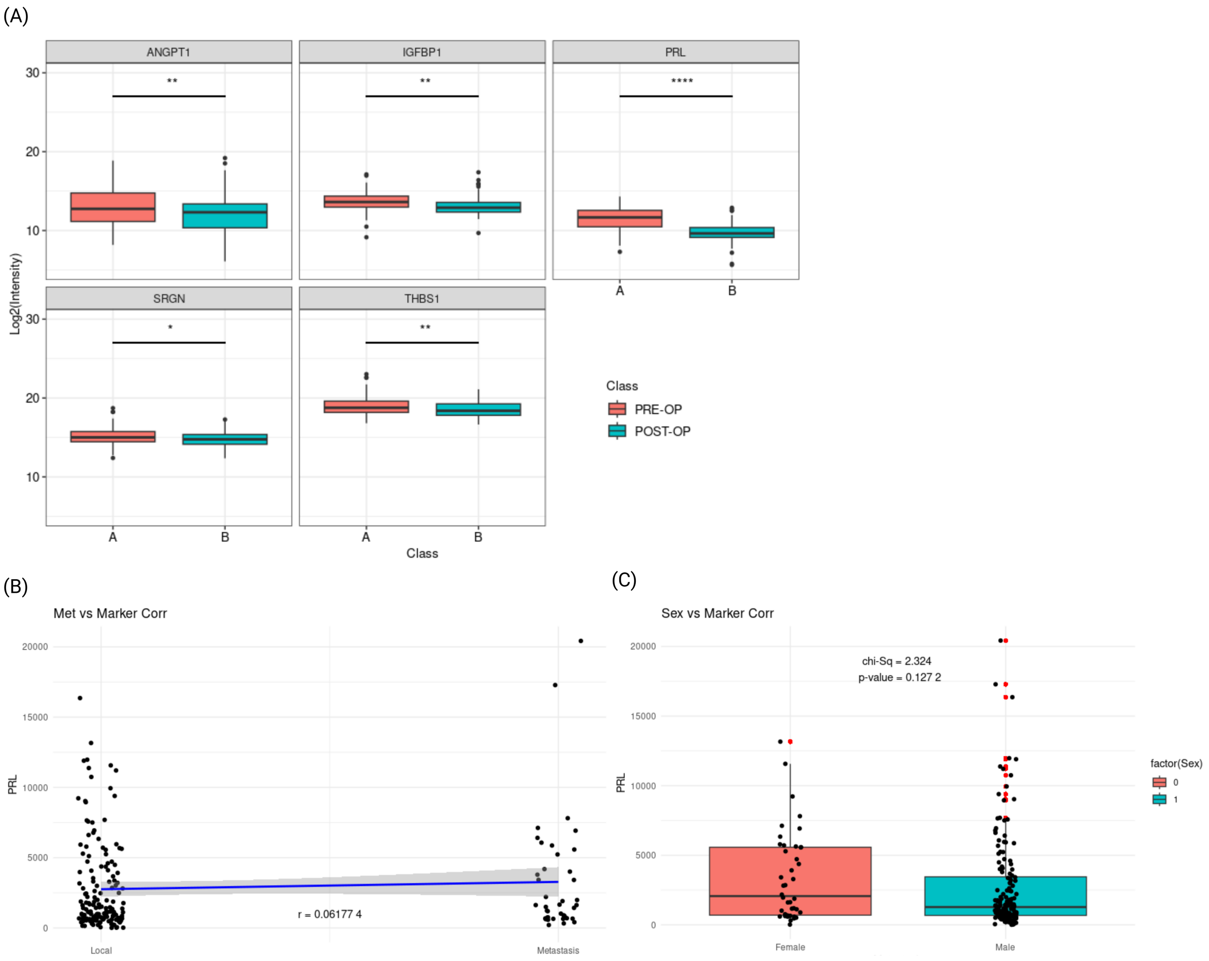


**Figure S3.** Defining signatures for localized ccRCC (A) Boxplot showing distribution of PRL, THBS1, ANGPT1, IGFBP1, SRGN between PRE-OP (orange) and POST-OP (blue) samples in Short-term FU Cohort (B) Scatter plot of observed lack of correlation of PRL in patients with localized ccRCC and recurrent mRCC (C) Scatter plot of observed lack of correlation of PRL in male and female ccRCC patients
